# Increased avidity for Dpp/BMP2 maintains the proliferation of eye progenitors in *Drosophila*

**DOI:** 10.1101/032995

**Authors:** M. Neto, F. Casares

**Affiliations:** CABD (Andalusian Centre for Developmental Biology). CSIC-UPO-JA. Campus Universidad Pablo de Olavide, 41013 Seville, Spain.; IBMC/Instituto de Investigaçâo e Inovaçâo em Saúde (i3S), Universidade do Porto. Rua Alfredo Allen, 208 4200–135 Porto, Portugal.

**Keywords:** *Drosophila* eye, progenitors, growth, Dpp/BMP2, *homothorax*, proteoglycans

## Abstract

During normal organ development, the progenitor cell state is transient: it depends on specific combinations of transcription factors and extracellular signals, that cooperate to sustain the proliferative potential and undifferentiated status of organ progenitor cells. Not surprisingly, abnormal maintenance of progenitor transcription factors may lead to tissue overgrowth, and the concurrence of specific signals from the local environment is often critical to trigger this overgrowth. Therefore, the identification of the specific combinations of transcription factors and signals that promote or oppose proliferation in progenitor cells is essential to understand normal development and disease. We have investigated this issue by asking what signals may promote the proliferation of eye progenitors in *Drosophila*. Two transcription factors, the MEIS1 homologue *homothorax (hth)* and the Zn-finger *teashirt (tsh)* are transiently expressed in eye progenitors causing the expansion of the progenitor pool. However, if their co-expression is maintained experimentally, cell proliferation continues and differentiation is halted. Here we show that Hth+Tsh-induced tissue overgrowth requires the BMP2 ligand Dpp and the activation of its pathway. In Hth+Tsh cells, the Dpp pathway is abnormally hyperactivated. Rather than using autocrine Dpp expression, Hth+Tsh cells increase their avidity for Dpp, produced locally, by upregulating extracellular matrix components. During normal development, Dpp represses hth and tsh ensuring that the progenitor state is transient. However, cells in which Hth+Tsh expression is maintained use Dpp to enhance their proliferation.

**Summary Statement:** In *Drosophila, homothorax*, the Meis1 homologue, and *teashirt* jointly sustain the proliferation of eye progenitor cells by increasing their avidity for BMP produced by the local microenvironment.

## Introduction

Normal organ growth depends on a balance between proliferation and differentiation of progenitor cells. Proliferation is controlled by intrinsic determinants (mostly transcription factors and epigenetic regulators) and extrinsic signals, acting locally or at longer range. In overproliferative diseases, specific aberrant combinations of intrinsic factors and signals result in deregulated growth and organ failure. Therefore it is essential to define these specific combinations of intrinsic and extrinsic factors to understand both normal development and overproliferative diseases. The development of the fly eye has been used to understand the mechanisms that control progenitor proliferation. In this system, the pax6-expressing progenitors are maintained undifferentiated and proliferative as long as they express a combination of two transcription factors: *homothorax (hth)*, a TALE-class homeodomain homologue to the MEIS1 proto-oncogene, and *teashirt (tsh)* a Zn-finger-type transcription factor also with homologues in mammals, the TSHZ gene family (Bessa et al., 2002). Importantly, when Hth+Tsh are forcibly maintained, progenitors remain proliferative and their differentiation into retinal cell types is blocked (Bessa et al., 2002; Peng et al., 2009; Lopes and Casares, 2010). Further studies showed that this proliferative effect was mediated, at least partly, through Hth and Tsh directly interacting with Yki, the YAP/TAZ homologue in the fly and transcriptional coactivator downstream of the Warts/Hippo (Wrts/Hpo) signaling pathway (Huang et al., 2005; Peng et al., 2009). In eye progenitors, Hth and Tsh would serve as transcriptional partners of Yki, with Hth providing a DNA-binding domain to the Hth:Tsh:Yki complex. The Wrts/Hpo pathway is a major controller of organ growth in flies and vertebrates and its malfunction has been also implicated in cancer in mammals (Dong et al., 2007). Wrts/Hpo may be in charge of translating information on tissue tension through a system that integrates planar epithelial polarity and cytoskeletal organization (Fernandez et al., 2011; Sansores-Garcia et al., 2011; Rauskolb et al., 2014). In addition to this global control, growth regulation is linked to organ patterning by long-acting signaling molecules. In the eye primordium, these signaling molecules (and their pathways) include wingless-Int (Wnt), Decapentaplegic (Dpp)/BMP2, Hedgehog (Hh) and JAK/STAT (Amore and Casares, 2010). However, neither the Wnt, JAK/STAT nor the Notch pathway (another non-autonomous but locally acting signaling pathway) seemed to contribute to the Hth+Tsh function (Peng et al., 2009). A potential role for Dpp has not been tested. The Dpp pathway had been shown to act antagonistically to *hth* and *tsh*, by repressing *hth* at long range and contributing to a shorter range repression of *tsh* as well (Bessa et al., 2002; Firth and Baker, 2009; Lopes and Casares, 2010). This, together with its potential antiproliferative role in the eye, made Dpp, in principle, an unlikely candidate to synergize with Hth+Tsh. However, Dpp is necessary for the proliferation of other discs, such as the wing (reviewed in (Restrepo et al., 2014)), and in the eye activation of the Dpp pathway can increase the proliferation of eye progenitors (Firth and Baker, 2009). In addition, while during normal development Dpp would ensure that high Dpp-signaling would be mutually exclusive with Hth+Tsh expression in progenitors, forcing Hth+Tsh maintenance might expose Hth+Tsh-expressing cells to high Dpp levels, which they would not encounter otherwise. Here we have tested specifically the effects that manipulating the Dpp/BMP2 pathway has on the maintenance of Hth+Tsh-driven progenitor state, as a model of progenitor-induced tissue overgrowth.

## Results and discussion

In the *Drosophila* eye, the fast proliferation of progenitors has been attributed to Hth and Tsh acting jointly as transcriptional cofactors of Yki in a cell-autonomous manner (Peng et al., 2009). Indeed, Yki overexpression alone also results in eye primordium overgrowths (Huang et al., 2005). However, the phenotypic outcome of Hth+Tsh and Yki-overexpressing eyes was different (Supplementary Figure 1). This different phenotype indicated that, although activation of the Hippo pathway contributes to the phenotype induced by Hth+Tsh, it cannot fully explain it, and pointed to the existence of additional inputs. To test the possibility that Dpp synergizes with Hth+Tsh in driving tissue overgrowth, we altered the expression levels of components of the Dpp pathway in the Hth+Tsh-overexpressing background (Figure 1 and Supplementary Figure 2). To evaluate the functional interactions, we analyzed changes in the size and extent of differentiation of late third instar eye primordia (also called imaginal discs), as well as the size and morphology of the eye primordium derivatives in adult flies. In general, manipulations that increased Dpp-signaling activity exacerbated the phenotype, while genotypes that weakened the pathway partly suppressed the Hth+Tsh phenotypes in discs and adult heads (Figure 1 and Supplementary Figure 2). The strongest interaction was observed with the type I Dpp receptor *thick veins (tkv):* RNAi-mediated silencing of *tkv (optix>hth+tsh+RNAi(tkv))* caused an partial rescue of the differentiated area in the eye primordium (as observed as Eya-expressing assembled ommatidia) and of the adult eye size (Figure 1C), whereas overexpression of an activated form of Tkv *(optix>hth+tsh +tkvQD)* dramatically enhanced the phenotype – no or very few ommatidia were detected in the eye primordium, which in addition was extensively overgrown and folded, and the adults showed large sacs of undifferentiated cuticle (Figure 1D).

**Figure 1.**
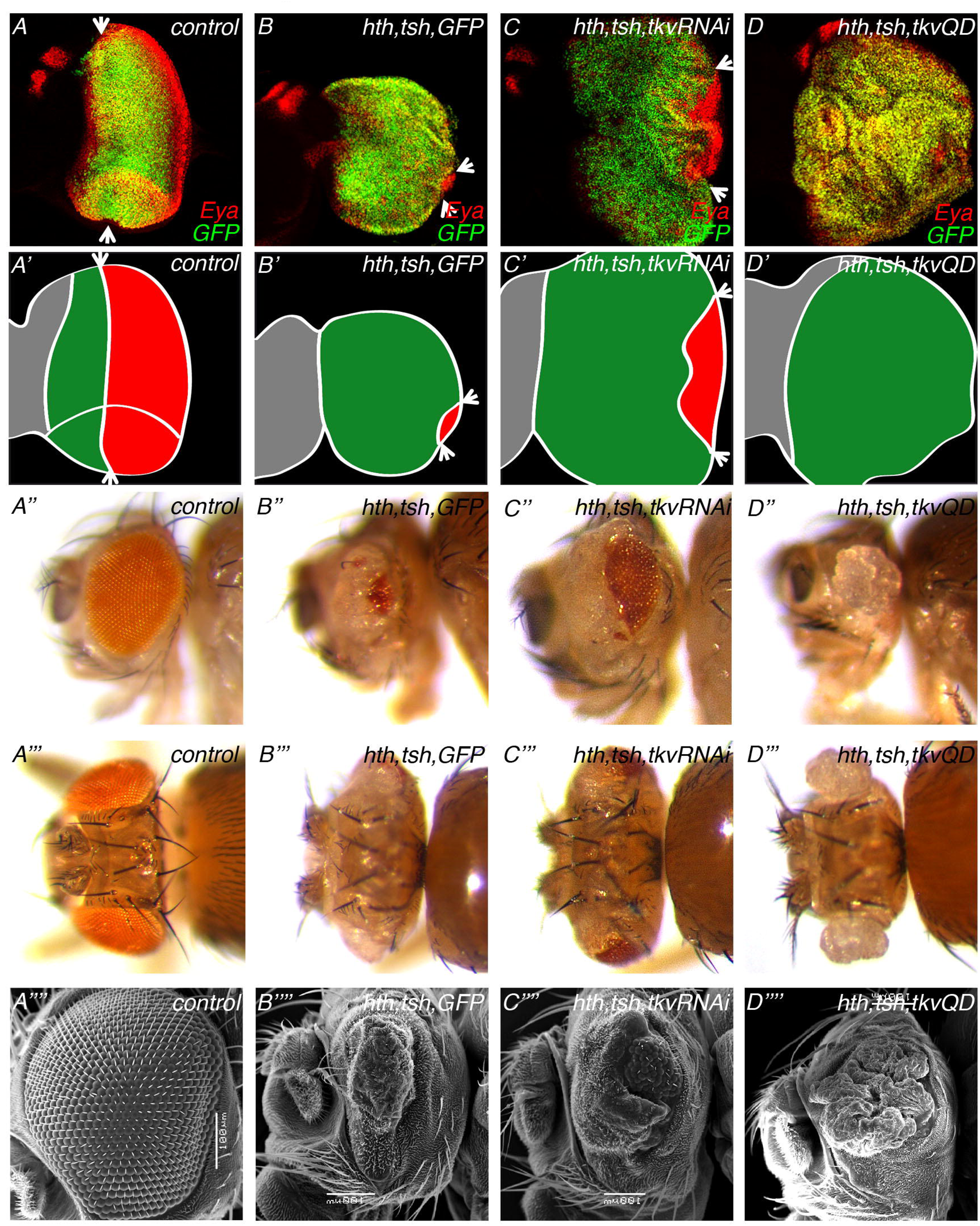
Altered expression of Tkv levels, either through RNAi or overexpression, in an Hth+Tsh background results in clear functional genetic interactions. Control late third instar (L3) eye disc (A) or eye discs expressing *Hth+Tsh+GFP* (B), *Hth+Tsh+tkvRNAi* (C) or *Hth+Tsh+tkvQD* (D) in undifferentiated cells using the optix2.3-GAL4 line. All discs are stained with anti-Eya (red); GFP in (A) comes from an UAS-GFP line, in (B) from UAS-GFP + UAS-131-GFPhth lines and in (C,D) from UAS-131-GFPhth line. Discs are oriented with anterior to the left and dorsal up (this orientation will be maintained throughout). Morphogenetic furrow is marked with arrows. (A’ - D’) Schematic representation of the discs above -antenna in grey, progenitor cells in green and differentiated cells in red. Lateral (A’’ - D’’) and dorsal views (A’’’ - D’’’) of adult heads of the same genotypes as above. (A’’’’ - D’’’’) SEM images of lateral views of adult heads of the above genotypes. In imaginal discs, Hth+Tsh overexpression resulted in the maintenance of progenitors and in the almost complete disappearance of the morphogenetic furrow, which marks the wavefront of retinal differentiation. Adult flies showed small patches of differentiated retina surrounded by an indistinct cuticle. Reducing the levels of Tkv by RNAi in an Hth+Tsh background resulted in a partial rescue of the morphogenetic furrow movement that led to a partly rescued eye. The expression of an activated version of Tkv (TkvQD) together with Hth+Tsh gave rise to highly folded discs with no morphogenetic furrow and overgrowths of indistinct cuticle in the adult.

We next analyzed the activity status of the Dpp pathway in Hth+Tsh cells by using phosphorylated-Mad (pMad), the active form of the nuclear transducer of the pathway, as a readout (reviewed by (Affolter and Basler, 2007). Of the three genotypes tested, only coexpression of Hth and Tsh *(optix>hth+tsh)* result in an increased pMad signal, affecting both its amplitude and range (Supplementary Figure 3). We note that in *optix>hth+tsh* discs, in which differentiation does not progress, the increased pMad signal is seen close to the margin, which is the normal site of production of *dpp before* differentiation onset. In order to determine whether the Dpp signaling increase is cell autonomous within the Hth+Tsh cells, and to more clearly analyze the spatial dependence of signal activation, we induced randomly located Hth+Tsh-expressing clones, and examined their effects on pMad levels, this time in both the wing and eye primordia. While cell clones expressing either Hth or Tsh alone did not show activation of the pathway (Figure 2A,B), some Hth+Tsh clones showed a strong, cell autonomous increase in pMad signal. Interestingly, only clones located near an endogenous source of Dpp (the AP boundary in the wing and the posterior margin and the morphogenetic furrow in the eye) exhibited high pMad, while pMad levels within the clones decreased as the clones were located farther away from these sources (Figure 2C,D). These results suggested that the activation of the Dpp signaling pathway in Hth+Tsh cells was induced by Dpp produced at its normal sites of production within the primordia, rather than by a Hth+Tsh-induced Dpp. To test this point specifically, we analyzed the expression of a *dpp* transcriptional reporter *(dpp-LacZ*, containing the “dpp-disc” enhancer (Masucci and Hoffmann, 1993) in Hth+Tsh clones in which pMad levels were increased. In these clones, *dpp-Z* was not activated (Supplementary Figure 4), ruling out the autocrine production of Dpp as a cause for Dpp-pathway activation. This fact, together with the requirement of the Dpp receptor *tkv* to promote the growth of Hth+Tsh tissue, led us to address directly the possibility that Hth+Tsh were using Dpp produced from local sources. To do so, we combined two binary gene expression systems: Gal4/UAS (Brand and Perrimon, 1993) and lexA/lexO (Yagi et al., 2010). A GFP-tagged form of Dpp (lexO-eGFP::Dpp) was driven using a *dpp-lexA* transgene. This driver was chosen for two reasons: first, it drove eGFP::Dpp in endogenous *dpp-* expression domains (i.e. in a stripe along the AP boundary in the wing primordium and around the posterior margin of the developing eye); and second, because the *dpp* enhancer (“dpp-disc” (Masucci et al., 1990; Muller and Basler, 2000)) used to drive lexA is not activated by Hth+Tsh (as shown in Supplementary Figure 4), effectively making the Dpp-producing system independent of Hth+Tsh. In this context, HA-tagged Hth+Tsh clones were randomly induced throughout the primordia. Indeed, Hth+Tsh-expressing clones retained high levels of eGFP::Dpp that were produced at *dpp-* expressing endogenous sites (Figure 3). We propose that maintenance of Hth+Tsh expression makes cells capture distantly produced Dpp that would then lead to the strong cell-autonomous pathway activation we observe. We noted that the levels of eGFP::Dpp in Hth+Tsh cells were high, suggesting an increased avidity for the Dpp ligand in these cells. Accumulation of eGFP::Dpp was detected even in clones that lie in regions where the surrounding eGFP::Dpp signal is low (Figure 3C’”). A similar phenomenon has been observed in clones expressing a membrane-tethered nanobody that traps eGFP (Harmansa et al., 2015).

**Figure 2.**
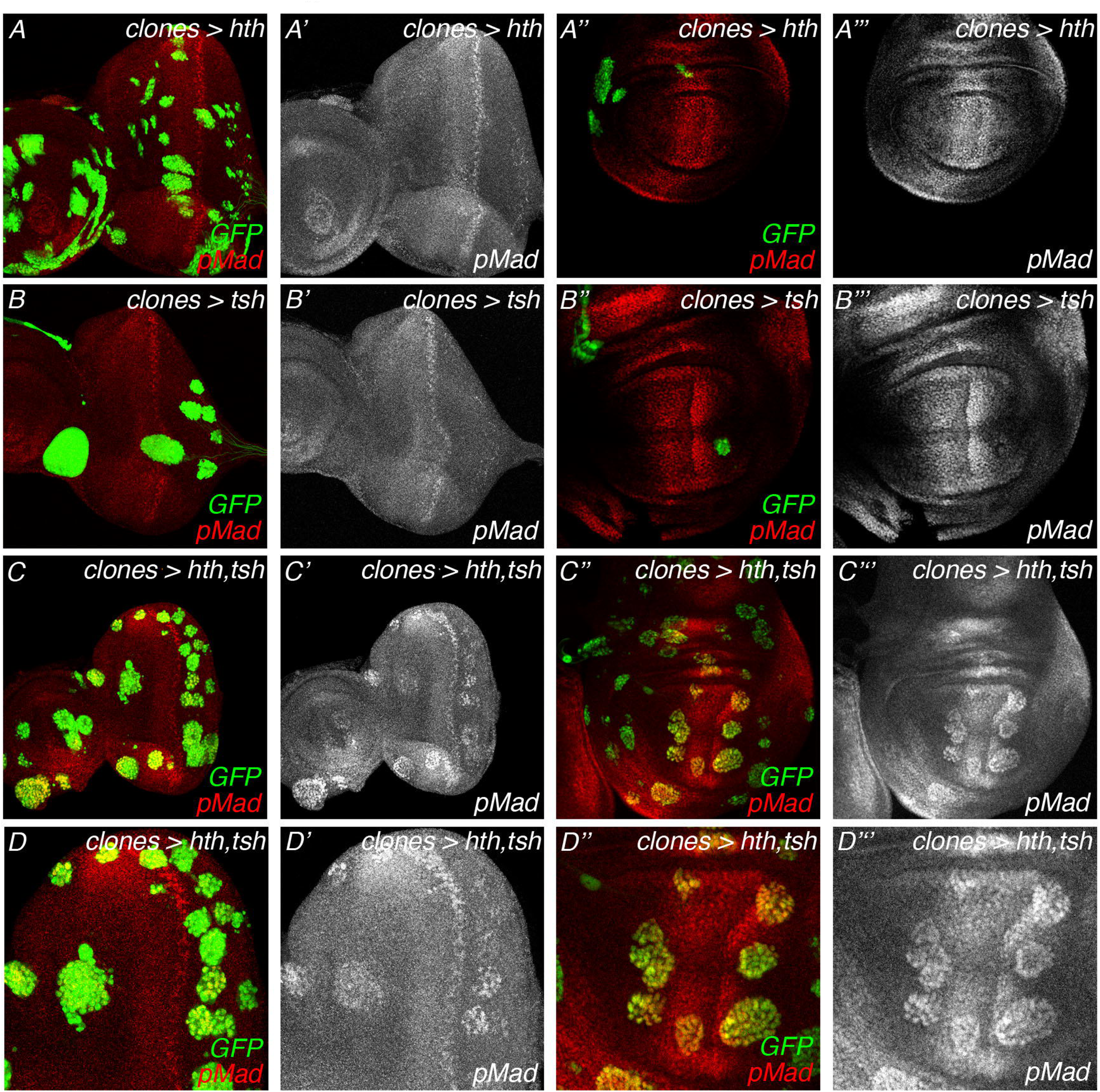
Forced maintenance of Hth+Tsh in clones results in a cell-autonomous accumulation of pMad. Hth- (A-A’’’), Tsh- (B-B’’’) or Hth+Tsh-expressing (C-D’’’) clones, marked by GFP, were induced in the eye and wing imaginal discs at 48–72 hours after egg laying. Anti-pSmad3 was used to detect endogenous pMad. Clones expressing Hth or Tsh alone did not show changes in the levels of pMad when compared to the wild-type neighboring cells. Hth+Tsh clones showed a spatial dependent accumulation of pMad in a cell-autonomous manner. pMad levels ranged from high in clones nearby sources of Dpp (AP boundary in the wing disc and posterior margin, and morphogenetic furrow in the eye disc) to background in clones located far away from these sources.

**Figure 3.**
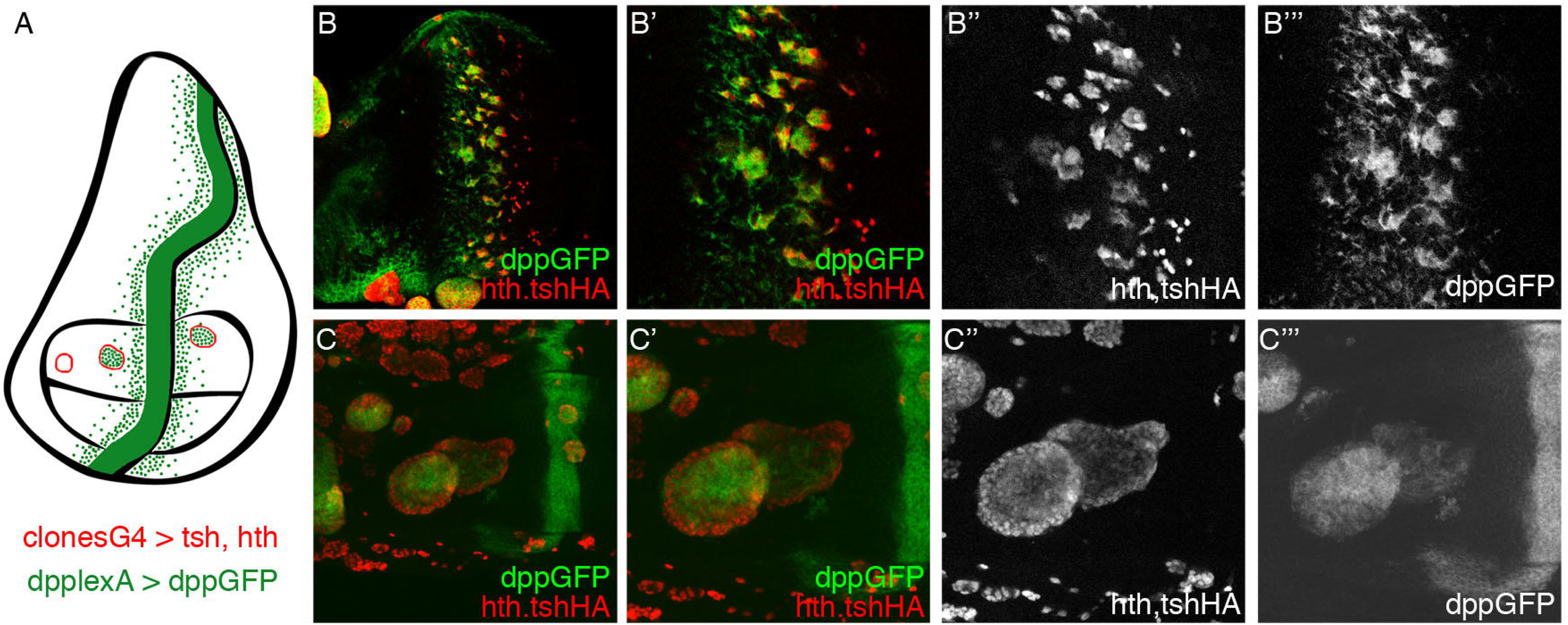
Hth+Tsh cells located near an endogenous source of Dpp are able to accumulate the morphogen. (A) Schematic representation of a Hth+Tsh wing imaginal disc using a combination of two binary gene expression systems. Green represents eGFP::Dpp produced at the endogenous dpp-expressing stripe along the AP boundary under the control of the lexA/lexO system. Hth+Tsh clones are represented in red, induced by the Gal4/UAS system. (B - C’’’) Hth+Tsh-expressing clones, marked by anti-HA, were induced in the eye and wing imaginal discs at 72–96 hours after egg laying using the Gal4/UAS system; simultaneously in the same discs eGFP::Dpp was expressed using the *dpp-lexA* driver through the lexA/lexO system. Hth+Tsh-expressing clones located near the Dpp source accumulated GFP-tagged Dpp.

One way to explain the increased avidity for Dpp would be if Hth+Tsh cells upregulated the expression of heparin-sulphate proteoglycans, extracellular matrix components known to act as key regulators of the distribution and activity of ligands such as Hh, Dpp or Wg (reviewed in (Yan and Lin, 2009). To test it, we analyzed the expression of the two glypicans *dally* and *dlp* in Hth+Tsh-expressing cells, using a *dally-Z* transcriptional reporter and an anti-Dlp antibody. Clones expressing Hth alone resulted in a cell autonomous downregulation of *dally* transcription, while the Dlp levels did not seem to be affected (Figure 4A-D). Tsh-expressing clones that fell within an endogenous Hth-expressing domain showed upregulation of *dally* transcription and Dlp levels, but all other clones did not (Figure 4E-H). However, Hth+Tsh clones showed high levels of *dally* transcription and Dlp membrane accumulation (Figure 4I-L). These results indicated that when cells maintain the progenitor transcription factors Hth *and* Tsh they increase the production of glypicans Dally and Dlp, facilitating the retention of endogenous Dpp. This retention would activate the Dpp-pathway and enhance Hth+Tsh-induced overgrowth and differentiation blockade.

**Figure 4.**
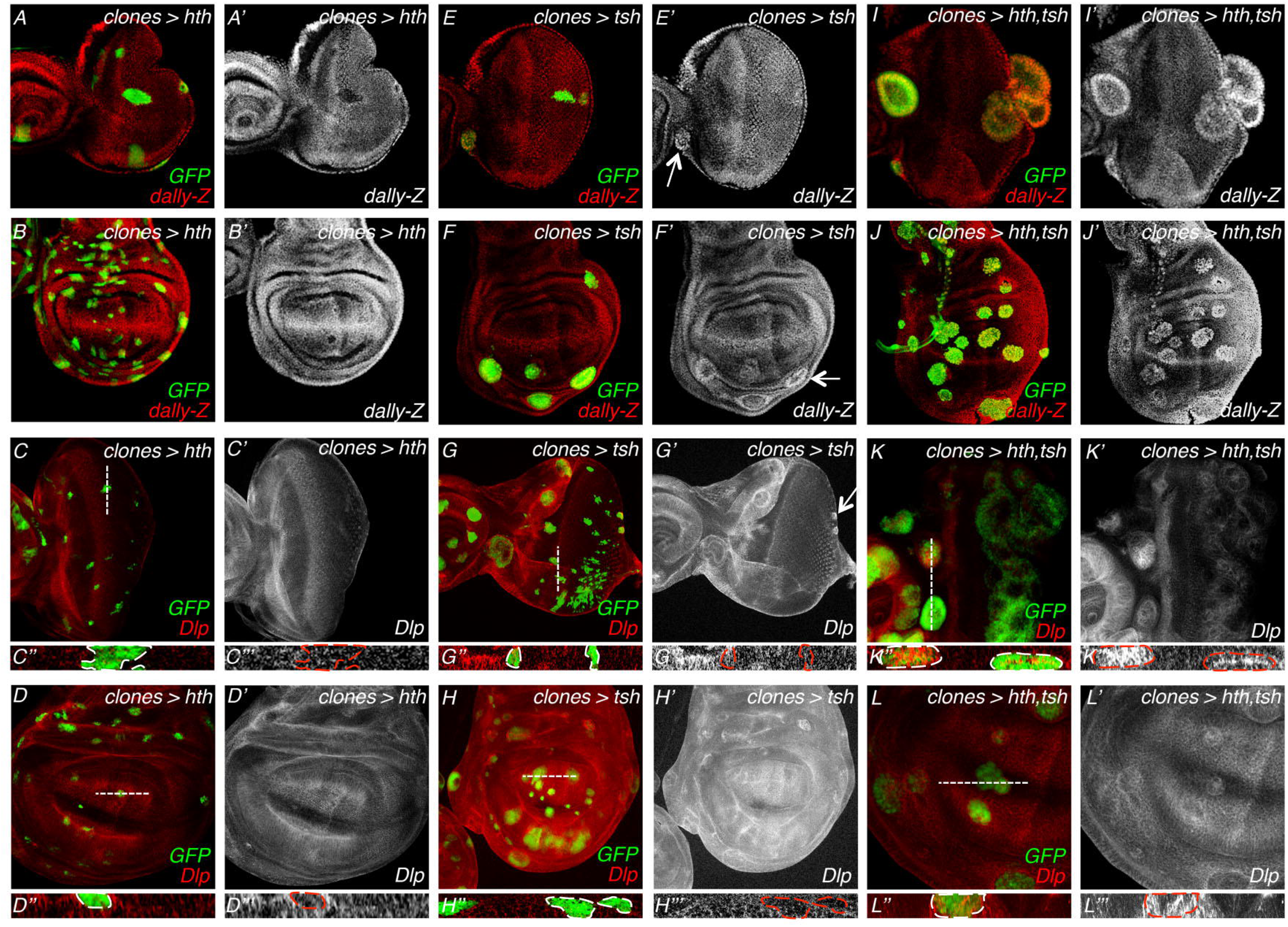
Hth+Tsh activate *dally* transcription and Dlp levels. Hth- (A-D’’’), Tsh- (E-H’’’) or Hth+Tsh-expressing (I-L’’’) clones, marked by GFP, induced in the eye and wing imaginal discs at 48–72 hours after egg laying. Clones induced in a dally-lacZ background were stained with anti-ß-galactosidase to monitor *dally* transcription (A-B’, E-F’, I-J’). The other discs (C-D’’’, G-H’’’, K-L’’’) were stained with anti-Dlp. The dashed lines approximately mark the optical cross-sections shown in (D’’, D’’’, H’’, H’’’, L’’, L’’’). Hth-expressing clones showed a decrease *dally-Z* levels and no significant changes in Dlp levels. Clones expressing Tsh did not show detectable differences in *dally* or Dlp, unless they were in a domain that expresses *hth* endogenously, in which case they showed higher levels of both glypicans (arrow). Hth+Tsh-expressing clones showed increased levels of *dally* transcription and Dlp protein.

Our results show that in the eye, like in the wing, Dpp also has the potential to promote proliferation. During normal eye development, though, high levels of Dpp signaling quickly repress *hth* (and at a shorter range also *tsh* (Bessa et al., 2002; Firth and Baker, 2009; Lopes and Casares, 2010)) and, by doing so, avoid the sustained exposure of progenitors to high signaling levels. However, experimental maintenance of Hth+Tsh reveals that Dpp can enhance proliferation of these Hth+Tsh expressing cells which, in addition, augment their avidity for Dpp, thus exacerbating the effect in the vicinity of Dpp sources.

## Materials and Methods

### Fly strains and genetic manipulations

All crosses were set up and raised at 25°C under standard conditions. The UAS/GAL4 system (Brand and Perrimon, 1993) and the lexA/lexO system (Yagi et al., 2010) were used for targeted misexpression. Fly stocks: *optix2.3-GAL4* (from R. Chen, Baylor College of Medicine); *UAS-yki* (Staley and Irvine, 2010); *UAS-GFP* (Bessa and Casares, 2005); *UAS-131-GFPhth* (Casares and Mann, 2000); *UAS-Flag-HA-tsh and UAS-Flag-HA-hth* (synthesized by C. M. Luque, Universidad Autónoma, Madrid, Spain); *yw*, hs-*FLP*^122^; act>y+>Gal4 (Struhl and Basler, 1993) with a recombined UAS-GFP transgene; *yw*, hs-FLP^122^, act>hsCD2>Gal4 (Basler and Struhl, 1994); *dpp-lacZ* (Masucci et al., 1990); *dpp-LHG86Fb* and *lexO-eGFP::Dpp* (Yagi et al., 2010) were a kind gift of K. Basler. dpp-pathway *lines: UAS-TkvRNAi* (VDRC #3059); *UAS-TkvQD* (Nellen et al., 1996); *UAS-puntRNAi* (VDRC #37279); *UAS-Punt* (Nellen et al., 1996); *UAS-Dad.T* (Tsuneizumi et al., 1997) and *UAS-Dpp.S* (Staehling-Hampton et al., 1994).

The dpp-pathway lines were crossed to the *optix2.3-GAL4,UAS-Flag-HA-tsh;UAS-131-GFPhth/SM6^TM6B* stock. Flies were observed under a LEICA MZ 9.5 stereomicroscope and pictures of adult heads from each genotype were taken with a LEICA DFC320 digital camera.

Random ectopic expression clones were generated using the flip-out method (Struhl and Basler, 1993). *yw, hs-FLP^122^; act>y+>Gal4;; UAS-GFP/TM6B* females were crossed to *UAS-Flag-HA-tsh, UAS-131-GFPhth* or *UAS-Flag-HA-tsh;UAS-131-GFPhth* males and transferred to 25°C. Clones were positively marked with GFP.

For the combination of Gal4/UAS and lexA/lexO systems, *yw, hs*-FLP^122^, *act>hsCD2>Gal4;UAS-Flag-HA-hth;lexO-eGFP::Dpp* females were crossed to *UAS-Flag-HA-tsh;dpp-LHG86Fb* males. Flip-out clones were induced by heat shock (10 minutes at 35,5°C) between 72h and 96h AEL and then maintained at 25°C. Clones were stained with anti-HA and anti-GFP.

### Immunostaining

Eye-antennal and wing imaginal discs from wandering third instar larvae were dissected and fixed according to standard protocols. Primary antibodies used were: mouse anti-Eya 10H6 at 1:100 (DSHB), rabbit anti-phosphoSmad3 1880–1 at 1:100 (Epitomics), used here to detect *Drosophila* phosphorylated-Mad (pMad) because its crossreactivity, mouse anti-ßGal at 1:1000 (Sigma), rabbit anti-ßGal at 1:1000 (Cappel), rat anti-HA at 1:500 (Roche), mouse anti-GFP at 1:1000 (Molecular Probes), mouse anti-Dlp 13G8 at 1:5 (DSHB), mouse anti-Armadillo N27A1 at 1:100 (DSHB). Alexa-Fluor conjugated secondary antibodies were from Molecular Probes. Images were obtained with the Leica SP2 confocal system and processed with Adobe Photoshop.

### Scanning electron microscopy (SEM)

Adult females were transferred to 75% ethanol for 24 hours at room temperature. Flies were dehydrated through an ethanol series (80%, 90%, 95% and twice 100%; 12–24 hours each step). Flies were then air-dried and mounted onto SEM stubs covered with carbon tape and sputter coated with gold (Edwards Six Sputter). Images were obtained using a JEOL 6460LV scanning electron microscope.

### pMad expression profiles

*optix>GFP, optix>131-GFPhth, optix>Flag-HA-tsh* and *optix>131-GFPhth+Flag-HA-tsh* L3 eye imaginal discs were stained simultaneously with anti-Arm and anti-pSmad3. Confocal imaging was done on the same day after the laser intensity had stabilized. The expression profiles were obtained using ImageJ. Signal intensity for anti-pSmad3 was measured in five independent discs. Measurements were taken ahead of the morphogenetic furrow, when present, or starting at the posterior margin of the disc when absent. The mean profile of each set of five profiles was represented in arbitrary units using Microsoft Excel.

## Acknowledgements

We thank K. Basler and the Bloomington and VDRC stock centers for fly stocks and the Developmental Studies Hybridoma Bank, University of Iowa, for antibodies, C.S. Lopes for critical reading of the manuscript, A. Iannini for technical assistance and the CABD Advanced Light Microscopy facility and the CITIUS (U. of Sevilla) SEM facility for imaging support.

## Author contributions

FC conceived the study; MN and FC designed the experiments; MN carried out the experiments; MN and FC wrote the manuscript.

## References

Affolter, M. and Basler, K. (2007) ‘The Decapentaplegic morphogen gradient: from pattern formation to growth regulation’, Nat Rev Genet 8(9): 663–74.

Amore, G. and Casares, F. (2010) ‘Size matters: the contribution of cell proliferation to the progression of the specification Drosophila eye gene regulatory network’, Dev Biol 344(2): 569–77.

Basler, K. and Struhl, G. (1994) ‘Compartment boundaries and the control of Drosophila limb pattern by hedgehog protein’, Nature 368(6468): 208–14.

Bessa, J. and Casares, F. (2005) ‘Restricted teashirt expression confers eye-specific responsiveness to Dpp and Wg signals during eye specification in Drosophila’, Development 132(22): 5011–20.

Bessa, J., Gebelein, B., Pichaud, F., Casares, F. and Mann, R. S. (2002) ‘Combinatorial control of Drosophila eye development by eyeless, homothorax, and teashirt’, Genes Dev 16(18): 2415–27.

Brand, A. H., and Perrimon, N. (1993) ‘Targeted gene expression as a means of altering cell fates and generating dominant phenotypes’, Development 118(2): 401–15.

Casares, F. and Mann, R. S. (2000) ‘A dual role for homothorax in inhibiting wing blade development and specifying proximal wing identities in Drosophila’, Development 127(7): 1499–508.

Dong, J., Feldmann, G., Huang, J., Wu, S., Zhang, N., Comerford, S. A., Gayyed, M. F., Anders, R. A., Maitra, A. and Pan, D. (2007) ‘Elucidation of a universal size-control mechanism in Drosophila and mammals’, Cell 130(6): 1120–33.

Fernandez, B. G., Gaspar, P., Bras-Pereira, C., Jezowska, B., Rebelo, S. R. and Janody, F. (2011) ‘Actin-Capping Protein and the Hippo pathway regulate F-actin and tissue growth in Drosophila’, Development 138(11): 2337–46.

Firth, L. C., and Baker, N. E. (2009) ‘Retinal determination genes as targets and possible effectors of extracellular signals’, Dev Biol 327(2): 366–75.

Harmansa, S., Hamaratoglu, F., Affolter, M. and Caussinus, E. (2015) ‘Dpp spreading is required for medial but not for lateral wing disc growth’, Nature.

Huang, J., Wu, S., Barrera, J., Matthews, K. and Pan, D. (2005) ‘The Hippo signaling pathway coordinately regulates cell proliferation and apoptosis by inactivating Yorkie, the Drosophila Homolog of YAP’, Cell 122(3): 421–34.

Lopes, C. S., and Casares, F. (2010) ‘hth maintains the pool of eye progenitors and its downregulation by Dpp and Hh couples retinal fate acquisition with cell cycle exit’, Dev Biol 339(1): 78–88.

Masucci, J. D., and Hoffmann, F. M. (1993) ‘Identification of two regions from the Drosophila decapentaplegic gene required for embryonic midgut development and larval viability’, Dev Biol 159(1): 276–87.

Masucci, J. D., Miltenberger, R. J. and Hoffmann, F. M. (1990) ‘Pattern-specific expression of the Drosophila decapentaplegic gene in imaginal disks is regulated by 3’ cis-regulatory elements’, Genes Dev 4(11): 2011–23.

Muller, B. and Basler, K. (2000) ‘The repressor and activator forms of Cubitus interruptus control Hedgehog target genes through common generic gli-binding sites’, Development 127(14): 2999–3007.

Nellen, D., Burke, R., Struhl, G. and Basler, K. (1996) ‘Direct and long-range action of a DPP morphogen gradient’, Cell 85(3): 357–68.

Peng, H. W., Slattery, M. and Mann, R. S. (2009) ‘Transcription factor choice in the Hippo signaling pathway: homothorax and yorkie regulation of the microRNA bantam in the progenitor domain of the Drosophila eye imaginal disc’, Genes Dev 23(19): 2307–19.

Rauskolb, C., Sun, S., Sun, G., Pan, Y. and Irvine, K. D. (2014) ‘Cytoskeletal tension inhibits Hippo signaling through an Ajuba-Warts complex’, Cell 158(1): 143–56.

Restrepo, S., Zartman, J. J. and Basler, K. (2014) ‘Coordination of patterning and growth by the morphogen DPP’, Curr Biol 24(6): R245–55.

Sansores-Garcia, L., Bossuyt, W., Wada, K., Yonemura, S., Tao, C., Sasaki, H. and Halder, G. (2011) ‘Modulating F-actin organization induces organ growth by affecting the Hippo pathway’, Embo J 30(12): 2325–35.

Staehling-Hampton, K., Jackson, P. D., Clark, M. J., Brand, A. H. and Hoffmann, F. M. (1994) ‘Specificity of bone morphogenetic protein-related factors: cell fate and gene expression changes in Drosophila embryos induced by decapentaplegic but not 60A’, Cell Growth Differ 5(6): 585–93.

Staley, B. K., and Irvine, K. D. (2010) ‘Warts and Yorkie mediate intestinal regeneration by influencing stem cell proliferation’, Curr Biol 20(17): 1580–7.

Struhl, G. and Basler, K. (1993) ‘Organizing activity of wingless protein in Drosophila’, Cell 72(4): 527–40.

Tsuneizumi, K., Nakayama, T., Kamoshida, Y., Kornberg, T. B., Christian, J. L. and Tabata, T. (1997) ‘Daughters against dpp modulates dpp organizing activity in Drosophila wing development’, Nature 389(6651): 627–31.

Yagi, R., Mayer, F. and Basler, K. (2010) ‘Refined LexA transactivators and their use in combination with the Drosophila Gal4 system’, Proc Natl Acad Sci U S A 107(37): 16166–71.

Yan, D. and Lin, X. (2009) ‘Shaping morphogen gradients by proteoglycans’, Cold Spring Harb Perspect Biol 1(3): a002493.

